# Aptamer charge-amplified field-effect transistor biosensors achieve picomolar detection limits for small-molecule biomarkers in complex biological matrices

**DOI:** 10.64898/2026.05.18.726065

**Authors:** Qitao Hu, Yasser Gidi, Hajime Fujita, Yihang Chen, Jenny Ji, Benjamin C. Wollant, Michael Eisenstein, H. Tom Soh

## Abstract

Aptamers are attractive receptors for small-molecule biomarker detection in complex samples because of their high stability, affinity, and specificity, but aptamer-based sensors generally lack the sensitivity to detect low-abundance analytes. As a solution, we developed the charge-amplified FET (CAFET) aptamer biosensor, which is designed to amplify the net charge variation within the Debye length that occurs as a consequence of aptamer-target binding. Our sensor utilizes a strand-displacement aptamer switch, which releases an initially-hybridized displacement strand (DS) upon target binding and thus induces a measurable net charge variation within the Debye length that is amplified to a large FET current response as signal readout. This signal can be further enhanced by adding a charge label to the DS. As a consequence, our sensor can achieve far greater sensitivity than previously described aptamer-FET sensors, where the binding-induced local charge variation is modest. We demonstrate 3-hydroxykynurenine and progesterone detection with a picomolar limit of detection in undiluted human plasma—four orders of magnitude lower than the dissociation constant (*K*_D_) of the aptamer component. The CAFET sensor design is modular and should be adaptable for the detection of a wide range of clinically-informative low-abundance analytes in complex samples.

## Introduction

The detection of small-molecule analytes in complex biological matrices is critical for biological research, clinical care, and personal health monitoring (1). Nucleic acid-based aptamers are particularly attractive bioreceptors for small-molecule biomarkers owing to their high stability, affinity, and specificity (2–4). However, many clinically-relevant small-molecule biomarkers have physiological concentrations in the nanomolar range or lower, and the applicability of aptamers for the detection of such low-abundance analytes is fundamentally restricted by their relatively high limit of detection (LOD). The detectable analyte concentration range for a typical aptamer biosensor is within two to three orders of magnitude of the aptamer’s dissociation constant (*K*_D_), assuming Langmuirian binding thermodynamics (5–7). Since most aptamers have a *K*_D_ in the high nanomolar or micromolar range (8), this typically results in a LOD in the low nanomolar range at best—beyond which the signal is overwhelmed by noise. This LOD limitation substantially impedes the aptamer-based detection of low-abundance biomarkers.

Field-effect transistor (FET)-based biosensors offer a promising solution owing to their inherent signal amplification capability (9–11). A typical FET device consists of three terminals—source (S), drain (D), and gate (G)—in which the current flowing from the source to the drain (*I*_SD_) is exponentially modulated by voltage applied to the gate (*V*_G_). The signal of the FET biosensor is based on charge variation within a short distance from the gate surface, known as the Debye length (*λ*_D_) (12, 13), where a change in *V*_G_ leads in turn to a much larger increase in the *I*_SD_ readout. In a typical FET-aptamer biosensor, the G electrode is functionalized with aptamer molecules that recognize an analyte of interest. Target binding induces a conformational change in the aptamer, which redistributes the native charges of the oligonucleotide backbone and thereby induces charge variation within the Debye length (14, 15). Early efforts in FET-aptamer sensor development have focused on small-molecule biomarker detection in buffer (16), but some recently developed FET-aptamer sensors are also compatible with biological samples, enabling neurotransmitter monitoring in brain tissue (17–19) and cortisol detection in sweat (20).

Despite these notable advances, FET-aptamer sensors have yet to fully deliver on their promise as high-sensitivity biosensors for detecting low-abundance small-molecule analytes in complex biological matrices like blood and plasma. For example, researchers have demonstrated detection in diluted serum, but could only reach micromolar detection limits which is not sufficient for many biomarkers that typically exist at nanomolar or picomolar concentrations (21). One major obstacle with traditional FET-aptamer sensors is the relatively small signal generated by the binding-induced charge redistribution of aptamer molecules within the Debye length. This is because the Debye length in physiological environments is extremely small (~0.7 nm)—shorter than the length of a trinucleotide sequence (22). This means that any aptamer conformational change is likely to involve only a small number of charges and have a commensurately small impact in terms of modulating the surface potential, and will thus fail to generate a sufficiently large signal for the sensitive detection of low-abundance analytes in complex biological matrices.

In this work, we present a charge-amplified FET (CAFET) aptamer biosensor that achieves a picomolar LOD in undiluted human plasma using aptamers with a modest, low-micromolar *K*_D_. CAFET uses a strand-displacement aptamer switch, which releases an initially-hybridized displacement strand (DS) upon target binding. The binding site for the DS is situated close to the electrode surface, and its displacement therefore induces a substantial net charge variation within the Debye length, producing a large current response as a signal readout. To further amplify the signal, we labelled the DS with a negatively-charged tag that resides within the Debye length, increasing the charge variation upon DS release while also exerting minimal effect on aptamer-target binding. Our approach produces a substantial net charge variation even at low target concentrations, and thus achieves greater sensitivity and superior LODs compared to traditional FET-aptamer sensors. To minimize the impact of biofouling from interferents in biological matrices and preserve CAFET’s sensitivity, we passivated the sensor surface with a polyethylene glycol (PEG) layer. As a demonstration, we have developed CAFET biosensors for 3-hydroxykynurenine (3HK) and progesterone, achieving picomolar LODs for both analytes in undiluted human plasma—four orders of magnitude lower than the *K*_D_ of the strand-displacement aptamers that we employed. These LODs are compatible with the detection of these analytes at physiological concentrations. It is worth noting that strand-displacement aptamers are a natural output of the well-established Capture-SELEX aptamer selection method, which has successfully delivered strand-displacement aptamers for various important biomarkers (23, 24). Therefore, we believe that CAFET should offer a generalizable platform for detecting low-abundance biomarkers in complex biological samples, even if the available aptamers exhibit affinities that are well above physiologically relevant concentration ranges for those analytes.

## Results

### Design principles of the CAFET biosensor

The fundamental limitation of conventional aptamer-based FET sensors is that they rely entirely on the charge redistribution of the aptamer molecule within the Debye length (**Fig. 1a**, upper), which inherently limits the magnitude of the *I*_SD_ response that they can generate. In our CAFET sensor design, on the other hand, the aptamer is hybridized to a complementary DS at a stretch of sequence adjacent to the sensor surface, such that several nucleotides from this strand penetrate the Debye region (**Fig. 1a**, lower). The target-induced release of the DS and its subsequent diffusion into the bulk solution alters the net charge within the Debye region, producing a substantial change in the FET surface potential that is then amplified to a huge *I*_SD_ response (**Fig. 1b**). The percent change in this metric is the principle readout for our sensor in this work. As discussed in more detail below, we can further enhance the magnitude of this readout by adding a charge label to the DS that also extends into the Debye region. These various signal enhancement effects collectively make it feasible to achieve LODs that surpass the lower bounds typically imposed by the aptamer *K*_D_ in conventional FET-aptamer sensors (**Fig. 1c**). Previous work has described a somewhat analogous FET sensor design that detects the target binding-induced charge release from a hairpin aptamer (25). However, this sensor could not operate in actual biological matrices, although it could work under diluted buffer conditions, since the aptamer was charge-labeled at sites in the stem region that would fall beyond the Debye region under physiological ionic conditions.

**Figure 1.**
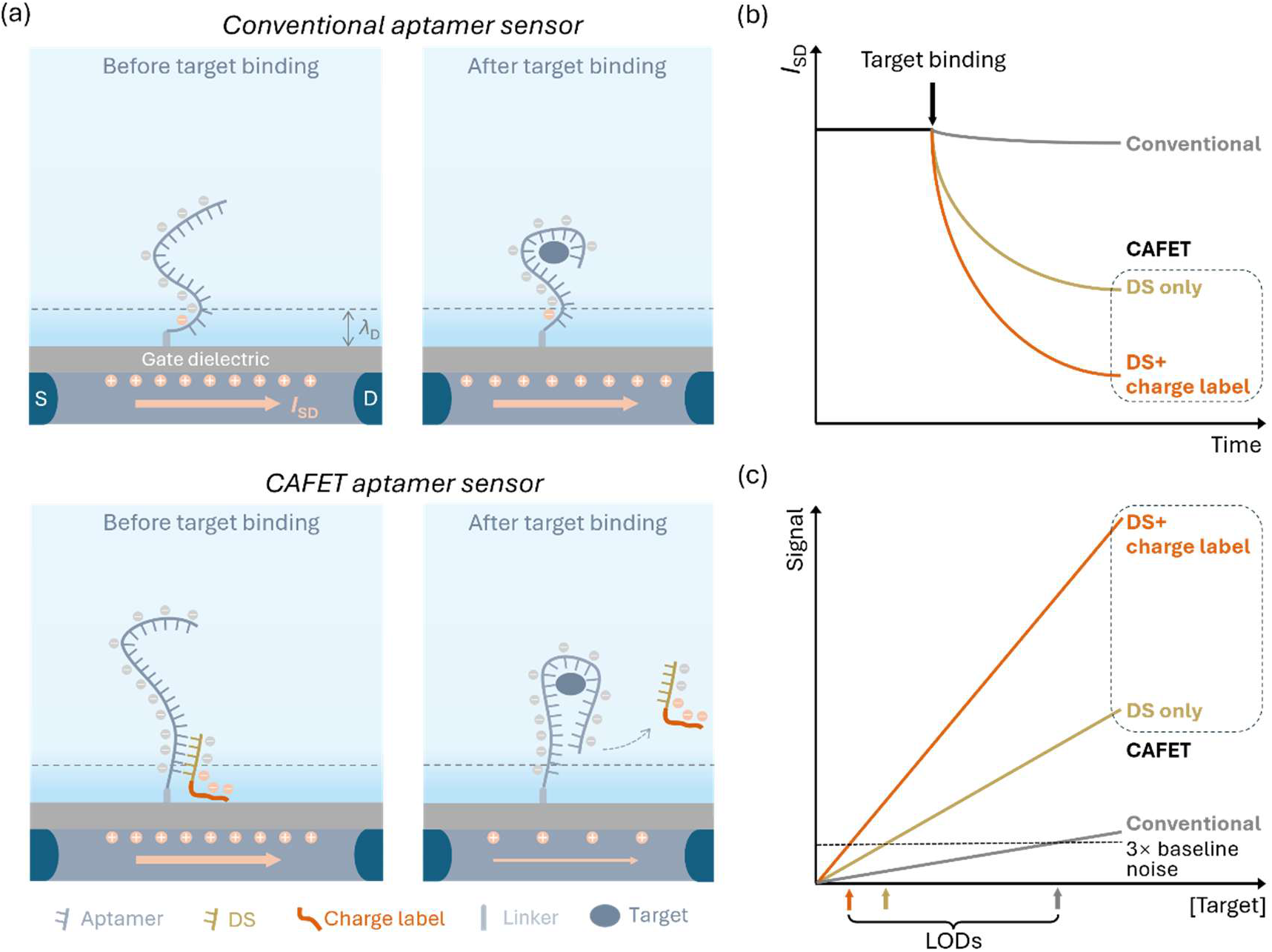
Design concept for our charge-amplified FET (CAFET) biosensor. (**a**) Conventional FET-aptamer sensors (top) rely entirely on the charge redistribution of aptamer molecules within the Debye length (*λ*_D_), resulting in minimal change in current *I*_SD_ in response to target binding. In our sensor design (bottom), the aptamer is initially hybridized to a complementary displacement strand (DS) that penetrates the Debye region. Target binding causes the aptamer to undergo a conformational change and release this DS sequence, leading to a substantial net charge variation (highlighted with negative charges) within the Debye length and thus a large change in *I*_SD_ as the DS dissociates into bulk solution. (**b**) Our CAFET sensor generates a greatly increased *I*_SD_ response compared to the conventional sensor. The magnitude of this response can be further enhanced by introducing a charge label onto the DS. (**c**) Illustration of how our CAFET sensor achieves a lower limit of detection (LOD).

### CAFET biosensor for detecting 3HK

We constructed our CAFET biosensor by functionalizing an extended-gate FET device with a strand-displacement aptamer (23). The extended-gate FET was fabricated by remotely connecting the G terminal of a commercially-available solid-state FET to an HfO_2_-coated Pt electrode (**Fig. 2a**; **Fig. S1**). We attached a circular polydimethylsiloxane (PDMS) barrier (diameter = 6 mm) onto the extended gate to serve as a sample chamber. The gate voltage was applied to the liquid sample via an Ag/AgCl reference electrode. As an initial demonstration, we constructed a FET sensor using a previously-reported 55-nucleotide (nt) strand-displacement aptamer that binds the kynurenine metabolite 3HK with a *K*_D_ of 6 μM (26). In our study, *K*_D_ was derived by fitting the sensor signal to the Hill equation (27, 28). Although the molecular system involves not only the aptamer-target interaction but also the DS dissociation, we have assumed a simplified equilibrium between the aptamer-target complex and the unbound state. Therefore, *K*_D_ represents an apparent value reflecting both the primary binding affinity and the downstream structural rearrangements (*i*.*e*., DS dissociation). We employed a 15-nt DS sequence that fully hybridizes to the 5’-terminal sequence of the aptamer, which is adjacent to the sensor surface (aptamer and DS sequence information is available in **Table S1** and **Fig. S2**). To produce a meaningful net charge variation within the Debye region, the aptamer nucleotides nearest to the sensor surface—and thus within the Debye region—must remain single-stranded once the DS has been released. We therefore ensured that the 7 nt at the 5’ end of the aptamer remain single-stranded even when the aptamer is fully folded. We developed a multilayered coating strategy to ensure high aptamer density on the extended gate. We first coated the surface with HfO_2_, and then chemically activated the HfO_2_-coated surface with piranha solution to generate hydroxyl groups, which we subsequently aminosilanized via treatment with (3-aminopropyl)triethoxysilane (APTES). The aminosilanized surface was then reacted with sulfosuccinimidyl (4-iodoacetyl)aminobenzoate (sulfo-SIAB). The 3HK aptamer was modified with a thiol group at its 5’ end, which enables covalent binding to sulfo-SIAB. Finally, we hybridized the immobilized aptamers to the DS via a thermal annealing process. The detailed immobilization protocol can be found in the Methods section and **Figure S3**.

**Figure 2.**
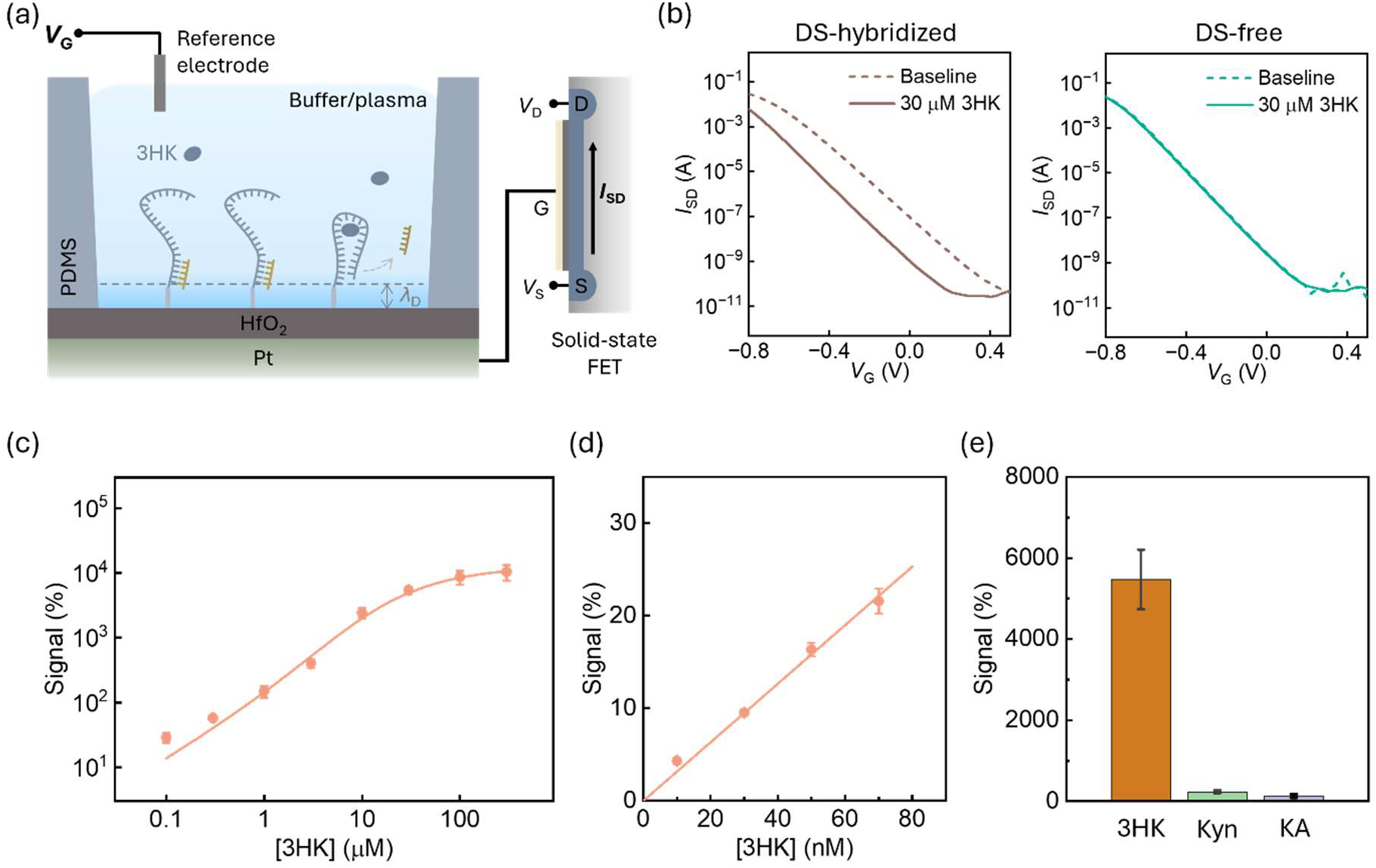
Design and construction of a FET-aptamer 3HK sensor. (**a**) Schematic of our extended-gate FET 3HK aptamer sensor. (**b**) Response to 30 μM 3HK by FET sensors functionalized with DS-hybridized (left) or DS-free (right) aptamers. (**c**) Binding curve for the DS-hybridized aptamer sensor versus 3HK. (**d**) FET signals in response to 0–70 nM 3HK. (**e**) Selectivity tests against 30 μM concentrations of 3HK or the structurally-similar molecules kynurenine (Kyn) and kynurenic acid (KA). Error bars in **c–e** were obtained from three devices.

To demonstrate our sensor’s response in high ionic strength conditions, we performed an initial analysis of 3HK in an aqueous buffer mirroring the ionic strength of undiluted plasma. After 1 hr incubation with 30 μM 3HK, we observed a clear leftward shift of the transfer curve, yielding a Δ*V*_G_ of 203 mV relative to our baseline measurement (**Fig. 2b**, left). Such a Δ*V*_G_ is transduced into a change of roughly two orders of magnitude in the FET current given the exponential *I*_SD_-*V*_G_ dependence. Real-time monitoring of the CAFET current in response to 30 μM 3HK revealed a Δ*I*_SD_ of roughly two orders of magnitude within 1 hr (**Fig. S4**), which is consistent with our estimate based on the Δ*V*_G_ measured in **Fig. 2b**. The real-time current trace also indicates that 1 hr target incubation is sufficient for stabilizing signals. To verify the function of the DS as a charged species that induced the signal change, we performed a control experiment with the DS-free 3HK aptamer, which generated a negligible response to target binding (**Fig. 2b**, right). We also confirmed that piranha pretreatment is crucial for DS release, as it makes the oxide surface more hydrophilic and thus suppresses hydrophobic interactions between the dissociated DS and the oxide surface, which could otherwise produce a false-positive response (**Fig. S5**).

We generated a binding curve from our CAFET sensor in response to a range of concentrations of 3HK (**Fig. 2c**). We obtained the signal data points by converting the original Δ*V*_G_ at various concentrations to percent current change based on the FET amplification effect. We then applied the Hill equation to the Δ*V*_G_ data, which was proportional to the number of aptamer molecules bound to the target, and converted it to a binding curve (**Fig. S6)**. In this fashion, we measured a *K*_D_ of 5.16 μM for our sensor, which is roughly equivalent to the originally reported value of 6 μM for this aptamer (26). We also determined the LOD of our sensor, which is defined as the analyte concentration at which the signal is three times greater than the standard deviation of the baseline. We extracted a sensitivity of 0.32%/nM at 3HK concentrations ranging from 0–70 nM (**Fig. 2d)**. Given the percent baseline deviation of 0.08% (**Fig. S7**), we determined that our CAFET sensor has a LOD of 730 pM—four orders of magnitude lower than the aptamer *K*_D_ (5.16 μM). Our sensor also demonstrated high selectivity against the structurally-similar molecules kynurenine (Kyn) and kynurenic acid (KA) (**Fig. 2e**). This high selectivity can be ascribed to the high specificity of the aptamer (26). After target binding and DS release, our CAFET biosensor can be regenerated by rehybridizing the DS to the aptamer (**Fig. S8**).

Our CAFET biosensor design delivered notable improvements in sensitivity relative to conventional FET-aptamer sensor designs, and we hypothesized that further gains should be achievable by incorporating a labeling strategy that increases the negative charge of the DS. To test this concept, we generated DS-(cT)_6_, a DS variant modified with negatively-charged 6-mer chains of oligo-carboxy-dT (cT) at its 3’ end. After hybridizing DS-(cT)_6_ to the aptamer, we observed a rightward shift in the transfer curve relative to the curve produced by the unmodified DS (**Fig. 3a**). We attribute this shift to the additional charges in the Debye region, as each cT 6-mer contributes two elementary charges. After 1 hr incubation with 30 μM 3HK, we observed a ~110 mV Δ*V*_G_ gain from the DS-(cT)_6_ sensor relative to the sensor with an unmodified DS; this in turn results in a 10-fold gain in *I*_SD_ (**Fig. 3b**). We confirmed that the magnitude of this signal enhancement is roughly proportional to the charges contributed by the labeled DS by testing DS variants labeled with shorter 2- or 4-mer cT chains, which generated a lower current signal gain than DS-(cT)_6_. We also demonstrated that our charge-labeling strategy works with other charged molecules such as oligo-dT and spacer C3, although these resulted in a lower signal gain since each unit of these charge labels contributes only one elementary charge (**Fig. S9**). We therefore used (cT)_6_ for our charge-labeled DS for all subsequent experiments.

**Figure 3.**
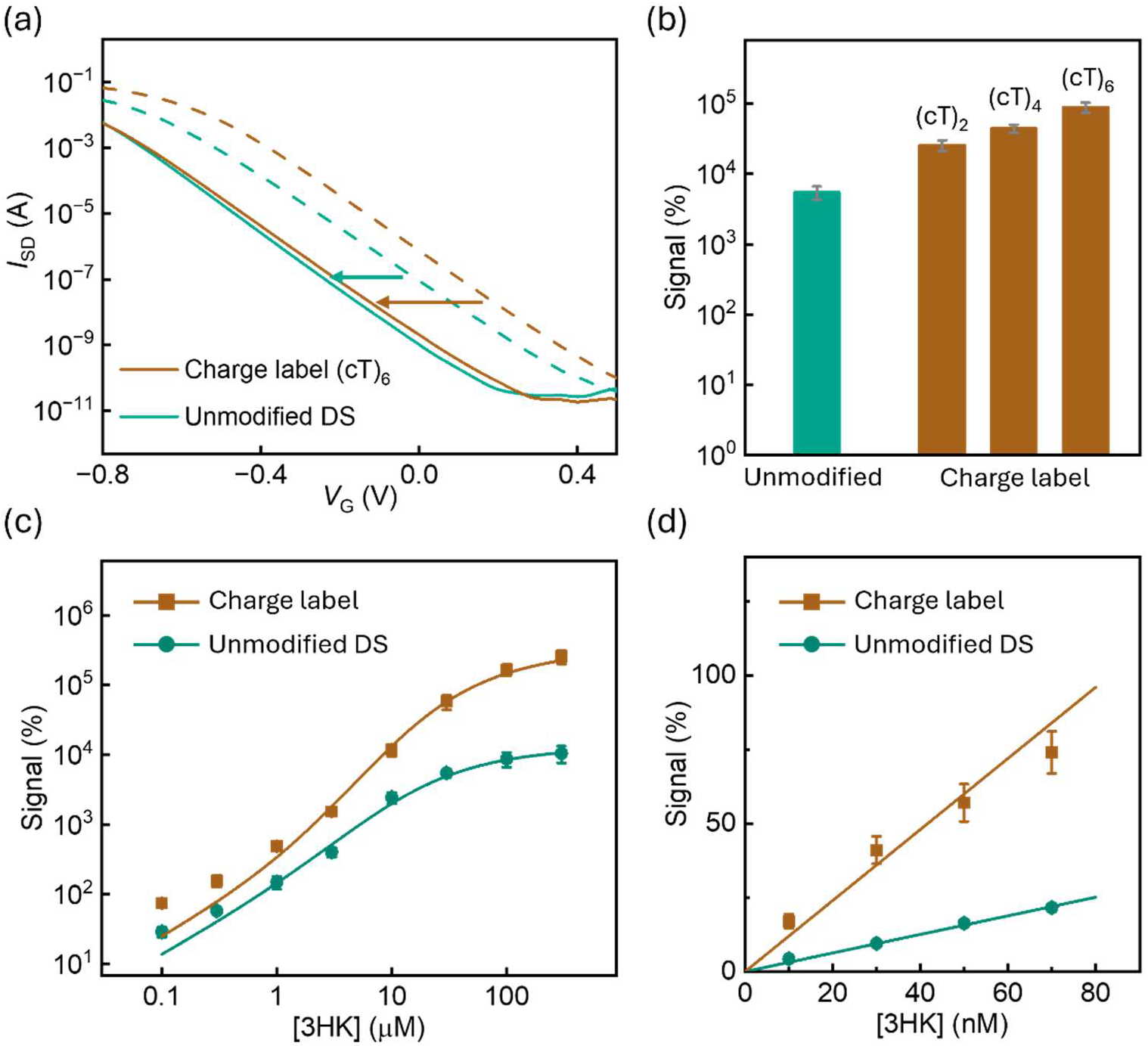
Charge-labeling further enhances CAFET sensor response. (**a**) Arrows depict the shifts in the FET transfer curve from baseline (dashed lines) after 1 hr incubation with 30 μM 3HK (solid lines) for sensors with unmodified DS or charge-labeled DS-(cT)_6_ sequences. (**b**) Signal response to 30 μM 3HK from CAFET sensors incorporating an unmodified DS or DS sequences modified with 2-, 4-, and 6-mer carboxy-dT (cT). (**c**) 3HK binding curves for sensors with unmodified or charge-labeled DS sequences. (**d**) Signal response to 0–70 nM 3HK from FET sensors with unmodified or charge-labeled DS sequences. All experiments were performed in high-ionic-strength buffer. Error bars in **b**–**d** were obtained from three devices.

We generated binding curves for 3HK using sensors with either the unmodified or charge-labeled DS, and confirmed that this modification does not meaningfully alter the affinity of the aptamer (*K*_D_ = 5.61 μM with charge-labeled DS versus 5.16 μM with the unmodified DS) (**Fig. 3c**). However, the charge-labeling strategy significantly enhanced the sensitivity of our sensor, yielding a LOD of 192 pM (**Fig. 3d**)—a notable improvement over the LOD achieved with the unmodified DS (730 pM). In addition, our sensor’s current response was notably higher compared to other previously-reported FET-aptamer biosensors operating under similar ionic conditions (**Table S2**). For example, at a target concentration equal to our aptamer’s *K*_D_, we realized a 5,238.9% current change with the charge-labeled DS sensor, whereas a previously-reported FET sensor based on a phenylalanine stem-loop aptamer showed only a 44.4% current signal at its *K*_D_ (25 nM) (21). Based on the signal gain with the charge-labeled DS relative to the unmodified one, we estimated the molecular density of aptamer to be 8 × 10^11^ cm^-2^ (**Fig. S10)** which is consistent with the literature (29, 30).

### CAFET detection in undiluted whole plasma

We next assessed how our CAFET sensors performed in the complex and interferent-rich molecular environment of undiluted pooled human plasma. We initially incubated our unmodified DS sensor with 30 μM 3HK in undiluted human plasma, and observed no measurable signal (**Fig. 4a**, ‘No PEG’). We attributed this result to the detrimental impact of biofouling, which physically obstructs target molecules from accessing the sensing surface (31), necessitating additional countermeasures to preserve sensor function in such complex samples. We therefore applied a PEG passivation strategy, which is widely utilized in the biosensor field (32). During sensor construction, we incubated various concentrations of thiol-modified PEG5000 and aptamer molecules with the sulfo-SIAB linker-bound HfO_2_-coated electrode surface. PEG passivation helped preserve the 3HK-specific sensor signal, with peak performance observed at 5% PEG (**Fig. 4a**). A further increase in PEG concentration to 10% produced a slight signal drop, which could be attributable to PEG competing for aptamer-binding sites on the sensor surface or compromising aptamer function due to steric effects (33). We also explored the passivation effect using PEG2000, and the resulting sensors exhibited a slightly lower signal compared to PEG5000 (**Fig. S11**).

**Figure 4.**
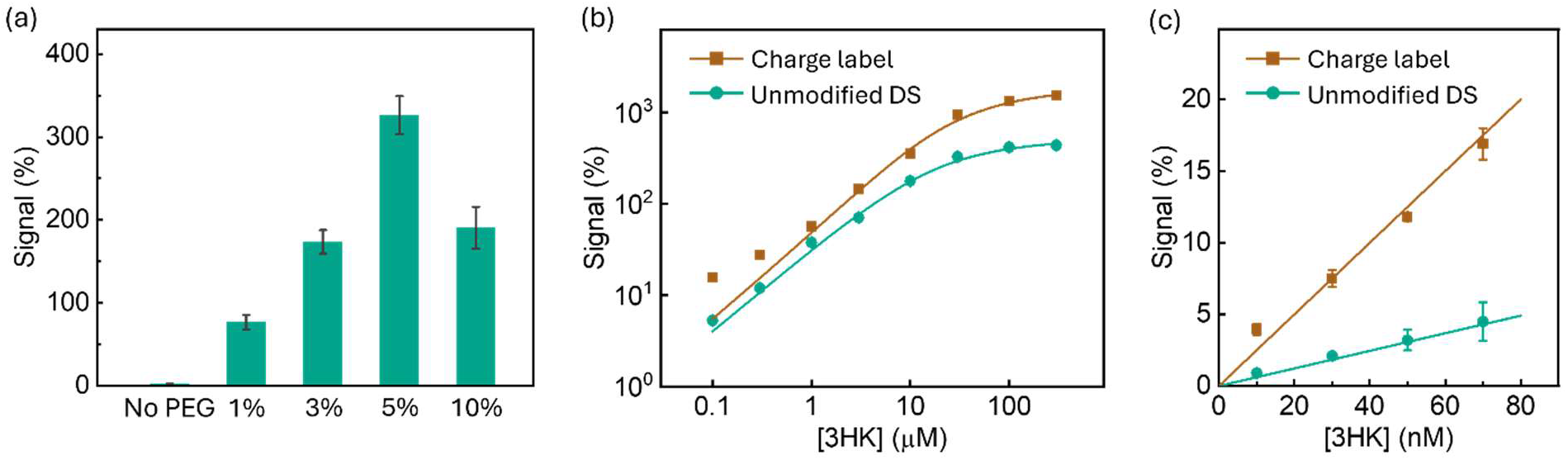
Detecting 3HK in undiluted human plasma. (**a**) Signal response from CAFET sensors passivated with varying molar fractions of PEG5000 to 30 μM 3HK in undiluted human plasma. The sensors were hybridized with unmodified DS and incubated with the target for 1 hr. (**b**) Binding curves for 3HK in undiluted human plasma from CAFET sensors passivated with 5% PEG5000 and hybridized with either unmodified or charge-labeled DS. (**c**) Signal from CAFET sensors with unmodified or charge-labeled DS in response to 0–70 nM 3HK in undiluted human plasma. Error bars were obtained from three devices.

We subsequently generated binding curves for 3HK in undiluted human plasma from sensors that were passivated with 5% PEG and hybridized to either unmodified or charge-labeled DS (**Fig. 4b**). The *K*_D_ values with both DS sequences were largely unchanged in plasma (unmodified DS: 6.72 μM; charge-labeled DS: 7.54 μM) compared to buffer (unmodified DS: 5.16 μM; charge-labeled DS: 5.61 μM). We measured a LOD of 840 pM for our charge-labeled DS sensor in undiluted plasma (**Fig. 4c**)—four times lower than what we observed with the unmodified DS (LOD = 3.4 nM)—confirming that our charge-labeling strategy leads to greatly improved analyte sensitivity in complex sample matrices. We also conducted sensing tests in 3HK-spiked plasma samples from three healthy donors (**Fig. S12**), and observed consistent responses in different plasma samples, validating the sensor’s accuracy with real-world specimens.

### Detecting clinically-relevant concentrations of progesterone in undiluted plasma

Finally, we demonstrated that our CAFET sensor platform can readily be adapted for the sensitive detection of other analytes. As an example, we selected the clinically important hormone biomarker progesterone, which plays crucial roles in the female menstrual cycle and reproductive function (34). Progesterone levels are also used as an indicator for conditions such as ectopic pregnancy, polycystic ovary syndrome, and ovarian cancer (35). We constructed a CAFET sensor using our recently-developed progesterone strand-displacement aptamer (36), which we combined with both unmodified and (cT)_6_ charge-labeled DS sequences. The design of the progesterone aptamer was similar to that of the 3HK aptamer, with a 15-nt DS sequence that fully hybridizes to the 5’-terminal sequence of the progesterone aptamer, which remains single-stranded once the DS has been released to ensure meaningful net charge variation in the Debye region (see **Table S1** and **Fig. S2** for sequence information). After generating binding curves for both sensors against a range of progesterone concentrations in buffer (**Fig. 5a**), we extracted *K*_D_ values of 4.83 and 1.22 μM for the unlabeled and charge-labeled DS sensors, respectively. These values correspond closely with previous measurements of this aptamer’s progesterone affinity (36). Our progesterone sensor also shows high selectivity against the structurally-similar interferents estradiol and cortisol (**Fig. S13**).

**Figure 5.**
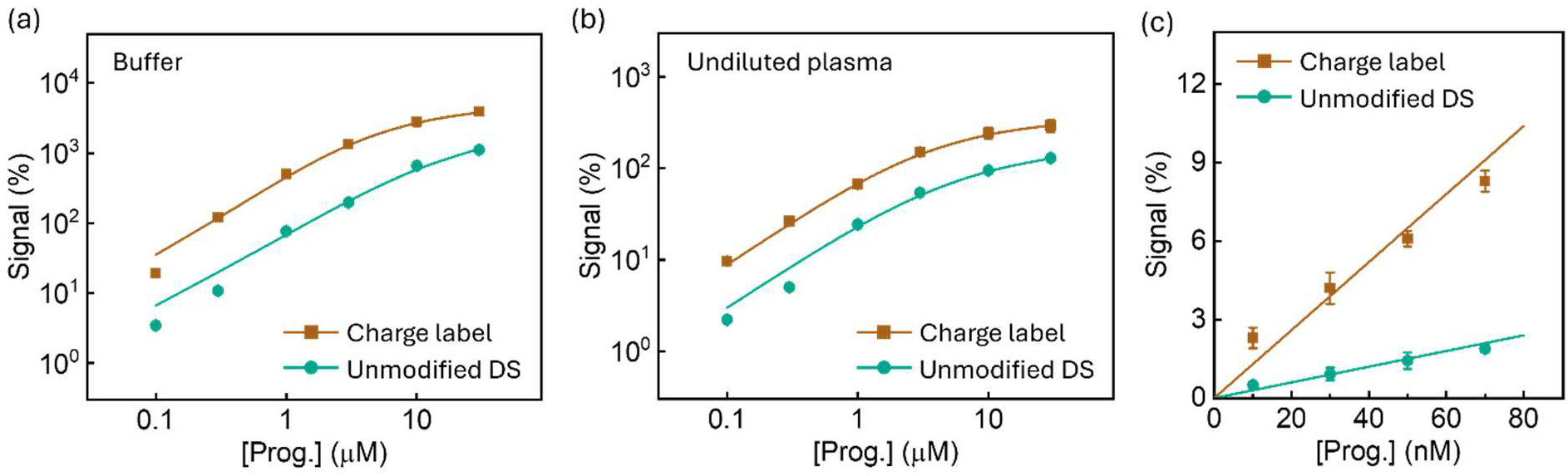
Adapting our FET-aptamer sensor design for progesterone detection. (**a, b**) Binding curves for progesterone in **a**, buffer and **b**, undiluted plasma from CAFET sensors hybridized with either unmodified or charge-labeled DS. The sensor used in the plasma experiment was passivated with 5% PEG5000. (**c**) Signal in response to 0–70 nM progesterone in undiluted human plasma by passivated CAFET sensors with unmodified or charge-labeled DS sequences. Error bars were obtained from three devices.

Importantly, this sensor could also detect progesterone in undiluted human plasma within a physiologically-relevant concentration range for pregnancy (**Fig. 5b**). For these experiments, we once again passivated the sensor with 5% PEG5000. The *K*_D_ values that we measured were largely unchanged in plasma (unmodified DS: 3.70 μM; charge-labeled DS: 1.78 μM) compared to buffer (unmodified DS: 4.83 μM; charge-labeled DS: 1.22 μM). We subsequently determined that our sensor achieves a LOD of 3.9 nM and 913 pM for progesterone in undiluted plasma with unmodified and charge-labeled DS sequences, respectively (**Fig. 5c**), again reflecting a LOD that was four orders of magnitude lower than the *K*_D_ of the aptamer. Given that typical physiological levels of plasma progesterone during pregnancy range from 23 to 730 nM, our sensor achieves the sensitivity and dynamic range that would be required to perform quantitative measurements of clinically-relevant concentrations of this analyte in patient samples. These results, together with those from our 3HK sensor, demonstrate that our CAFET sensor design is well-suited for achieving sensitive detection of a variety of analytes in complex samples with minimal interference due to biofouling.

## Discussion

In this work, we present CAFET—a novel FET-aptamer sensor platform that achieves LODs far below the *K*_D_ of their aptamer component in complex biological samples. In contrast to conventional sensing mechanisms that rely on aptamer charge redistribution, we leverage a DS release event that generates a large net charge variation within the Debye length. Our aptamer switches are initially hybridized to a DS such that the negative charge of this complementary strand extends into the Debye region. When the target binds the aptamer, the resulting conformation change leads to the release of the DS, producing a sizable modulation of the charge within the Debye region; this shift in voltage is subsequently transduced into an exponential change in output current. The magnitude of this response can be further enhanced via the addition of charge-labeling groups to the DS, which increases the sensitivity of the sensor by several fold. We also incorporated a PEG passivation process that effectively eliminates the problem of biofouling, which is a persistent issue with complex biological samples. We developed CAFET sensors for 3HK and progesterone, and in both cases achieved picomolar LOD values in undiluted human plasma, enabling detection of these analytes at physiologically-relevant concentration ranges. We believe that this same sensor design strategy should be broadly generalizable, given that target-specific strand-displacement aptamers can readily be isolated for various biomolecules via the widely-used capture-SELEX process (23, 24). Given the signal amplification mechanism demonstrated in this work, even an aptamer switch with relatively modest affinity could be expected to deliver robust sensing performance when incorporated into our CAFET framework. The LOD of our sensors can be further decreased by hybridizing shorter DSs to the aptamers, which results in a lower *K*_D_ due to the greater ease with which the DS can be displaced by target binding. For instance, in previous work we achieved a ~10-fold reduction in the *K*_D_ of our progesterone aptamer by replacing a 15-mer DS with a 13-mer (36). Importantly, the CAFET design should also be applicable to other sensor substrates, as the HfO_2_ gate dielectric layer upon which the aptamer molecules were immobilized is also compatible with other common FET materials such as Si (37), carbon nanotubes (38), and graphene (39). The extended-gate FET configuration of our sensors is also compatible with flexible and biocompatible substrates (40), making this design a potentially good fit for implementation in contexts such as wearable biosensors.

## Materials and Methods

### Materials

All aptamer and DS sequences were chemically synthesized and high-performance liquid chromatography (HPLC)-purified in our lab (sequences are shown in **Table S1**). 3-hydroxykynurenine (3HK) (#H1771), progesterone (#P8783), (3-aminopropyl)triethoxysilane (APTES) (#440140), Na_2_CO_3_ (#223530), and NaHCO_3_ (#S5761) were ordered from Sigma-Aldrich. Acetone (#A929-4), sulfosuccinimidyl (4-iodoacetyl)aminobenzoate (sulfo-SIAB) (#PI22327), and Tris(2-carboxyethyl)phosphine (TCEP) (#20490) were purchased from Thermo Fisher Scientific. Phosphate-buffered saline (PBS, pH 7.4, 1×) buffer was ordered from Gibco (#10010023). Thiol-modified PEG (5K and 2K) was purchased from Laysan Bio (#176-53 and 176-159). The high-salt buffer consists of 20 mM Tris-HCl, 120 mM NaCl, 5 mM KCl, 1 mM MgCl_2_, 1 mM CaCl_2_, and 0.01% Tween-20 in nuclease-free water.

### Extended-gate FET fabrication

The extended-gate FET was constructed by connecting the gate region of the commercial p-type FET DigiKey SIA517DJ-T1-GE3 and an extended-gate electrode fabricated in the Stanford Nanofabrication Facility (SNF) cleanroom using CMOS-compatible processes. The substrate of the gate electrode was a standard 4” bulk Si wafer (p-type boron-doped, [100], 0.1–0.9 Ω‧cm, 500-μm thickness, single-side polished; NanoSilicon Inc.). A 100-nm-thick Pt layer was deposited on top of the Si wafer with a 10-nm-thick Ti adhesion layer via electron beam evaporator (ATC-E, AJA International). Pt and Ti were deposited at a rate of 1 Å/s and a base pressure of 1 × 10^-7^ Torr. The substrate was kept at room temperature without additional heating. Then a 5-nm-thick HfO_2_ layer was coated onto the Pt layer via atomic layer deposition (Fiji, Veeco Instruments) using a plasma-enhanced mode at 200 °C. The precursor for HfO_2_ is Tetrakis(dimethylamido)hafnium(IV) ([(CH_3_)_2_N]_4_Hf). The pulse and purge times are both 5 s. The 5 nm thickness was reached within 50 cycles. The 4” wafer was then cut into 2-cm × 3-cm pieces for use as extended-gate electrodes.

### Aptamer immobilization

The extended-gate electrode was pretreated using piranha (9:1 sulfuric acid:hydrogen peroxide, v/v) at 120 °C for 15 min. After cleaning in DI water and drying with N_2_ flow, the electrode surface was aminosilanized in a mixture of 150 μL APTES and 10 mL acetone for 5 min at room temperature. The surface amino groups were then bound to sulfo-SIAB by incubating the electrode in 10 mM sulfo-SIAB in 100 mM NaHCO_3_ (pH 8.3) for 40 min at room temperature. The electrode was then incubated in 2 μM thiol-modified aptamer solution in 100 mM NaHCO_3_/Na_2_CO_3_ buffer (v/v = 92:8, pH 9.2) for 40 min at room temperature. Finally, the electrode was annealed with 10 μM DS in high-salt buffer at 90 °C for 5 min and then allowed to equilibrate at room temperature for 40 min. For PEG surface passivation, thiol-modified PEG molecule was mixed with the thiol-modified aptamer at a molar ratio of 1, 3, 5, or 10% during the aptamer incubation step. The thiol-modified aptamer and PEG were pretreated with 1,000× concentrated TCEP solution in 1× PBS buffer for 1.5 hrs at room temperature in the dark.

### Liquid measurements

We used a differential measurement setup to obtain our biosensing signals to calibrate the baseline drift. A pair of 6-mm-diameter PDMS chambers were attached onto the extended gate electrode; one held 100 μL of analyte-free background solution (baseline) while the other held 100 μL of analyte solution for synchronous incubation (**Fig. S1**). To ensure strong adhesion to the gate electrode, the PDMS container was processed in ozone for 3 min before attachment. FET measurements were performed using a Semiconductor Analyzer B1500A (Keysight Technologies). The gate voltage was applied to an external Ag/AgCl reference electrode (EP2, World Precision Instruments) inserted into the baseline or analyte chambers. FET transfer curves were obtained by monitoring *I*_SD_ while sweeping the gate voltage. All transfer curves were characterized while fixing the drain voltage at 1 V and at room temperature.

## Acknowledgments

We are grateful for financial support from the Helmsley Charitable Trust and Wellcome Leap SAVE program. Q.H. acknowledges the Wallenberg Foundation Postdoctoral Scholarship (KAW 2021.0334). We thank Nghi Torres for her assistance with custom DNA synthesis.

## Author Contributions

Q.H. and H.T.S. conceived the project. H.T.S. supervised the project. Q.H. initiated the charge labelling idea. Q.H., Y.G., and H.T.S. designed the experiments. Q.H. performed the experiments and collected and analyzed the data. Y.G. advised on the surface treatment and the aptamer design and immobilization. H.F. advised the progesterone aptamer design. Q.H., H.F., and J.J. discussed the binding model. Q.H. and H.T.S. co-wrote the paper. All authors discussed the results and commented on the paper.

## Data availability

The data that support the findings of this study are available within the paper and its Supplementary Information files.

