## Supplemental Information for "Aptamer charge-amplified field-effect transistor biosensors achieve picomolar detection limits for small-molecule biomarkers in complex biological matrices"

This PDF file includes:
Figure S1 to S13
Table S1 and S2

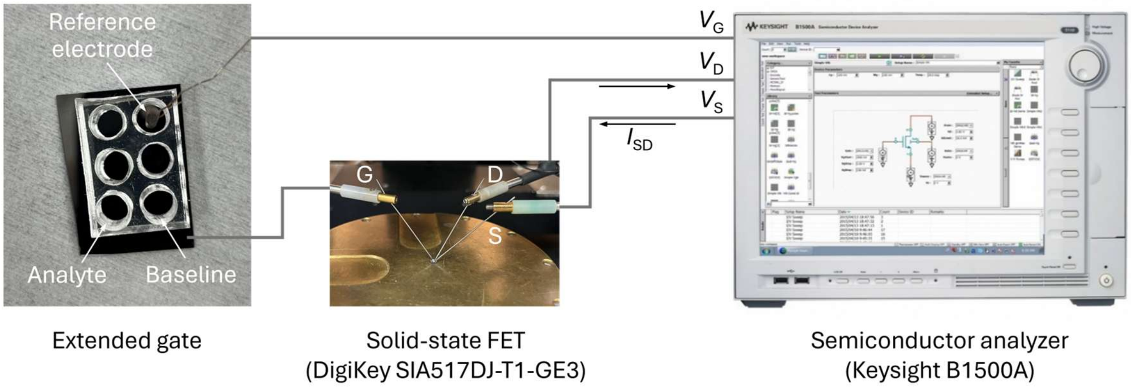


**Figure S1. Measurement setup of extended-gate FET biosensors.** We used a differential extended gate configuration to collect the analyte signals. The extended gate ($HfO_2$/Pt/Si substrate) was separated into 2 × 3 regions by PDMS containers. All six regions were functionalized with the same aptamer molecules in a single immobilization batch; the left three were incubated with analyte solution, while the right three were incubated with analyte-free solution to measure the baseline. We used a standard semiconductor analyzer (Keysight B1500A) to measure transfer curves of the extended-gate transistor. The gate voltage $V_G$ was applied via an Ag/AgCl reference electrode to the extended gate, which was connected to the gate terminal of a commercial solid-state transistor (DigiKey SIA517DJ-T1-GE3) using a probe station. The source-to-drain current ($I_{SD}$) of the transistor was monitored by the semiconductor analyzer while sweeping $V_G$. The sensing signal was extracted as the transfer curve shift after analyte incubation relative to the baseline. The signals of the three analyte-background pairs on one extended electrode were used to generate the average signals and error bars.

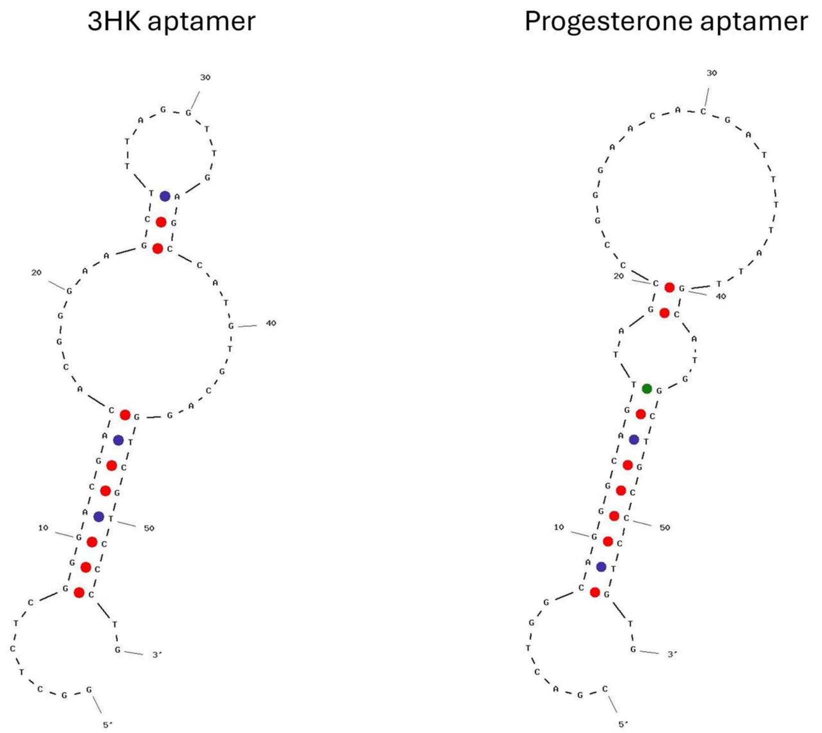


**Figure S2. Secondary structures of the aptamers for 3HK and progesterone**. The 3HK and progesterone aptamers have the same length (55 nt). The blue and red dots indicate A-T and C-G base-pairs, respectively; green is the wobble base pairing of G-T.

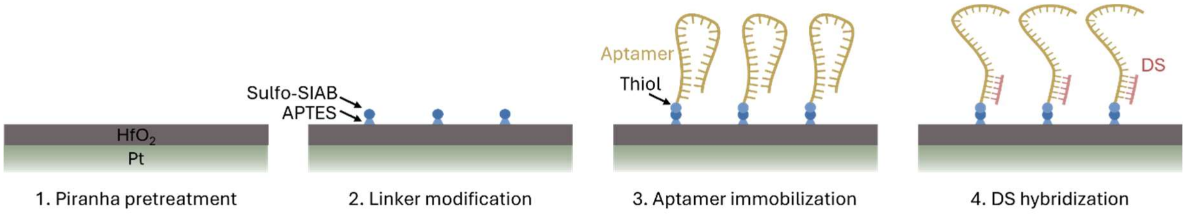


**Figure S3. Schematic illustration of aptamer immobilization steps on the extended-gate electrode.**

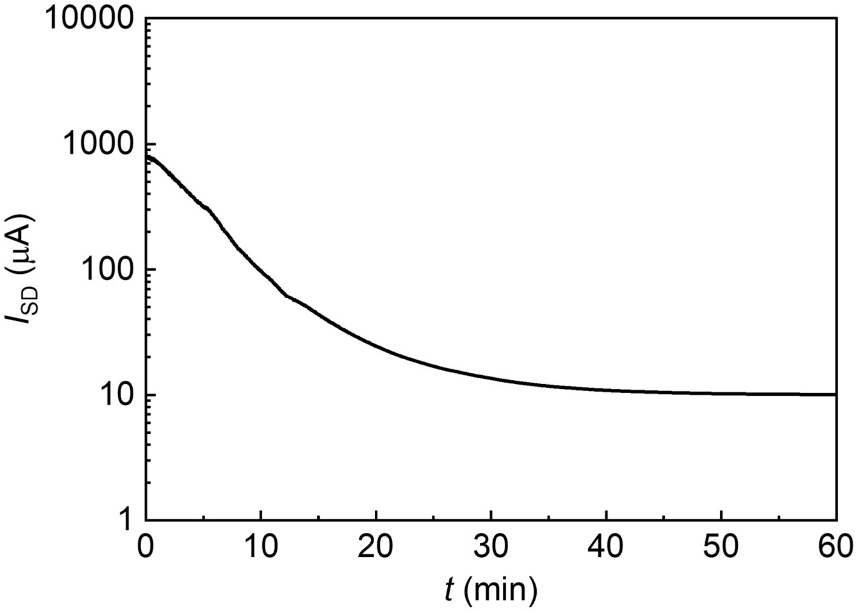

**Figure S4. Time response curve to 30 μM 3HK.** The 3HK sensor was hybridized to the unmodified DS. The data were obtained in high salt buffer.

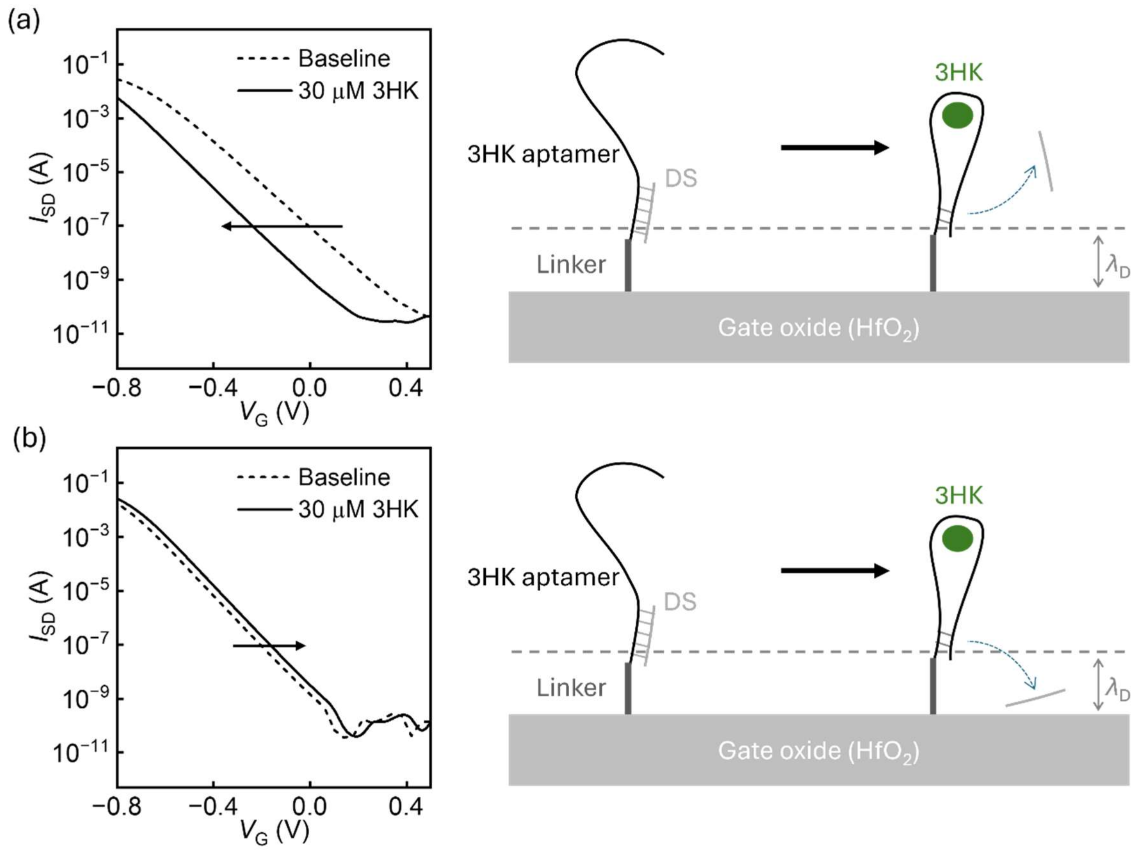


**Figure S5. Assessing the importance of piranha pretreatment.** Transfer curve shifts from FET-aptamer 3HK sensors prepared (**a**) with and (**b**) without piranha pretreatment before aptamer immobilization. The transfer curve (left) shifts to the opposite polarity without piranha treatment compared to the device prepared with piranha treatment. This could be due to the adsorption of the de-hybridized DS onto the oxide surface via hydrophobic interaction (right), which introduces additional negative charges into the Debye region. In contrast, piranha treatment makes the oxide surface more hydrophilic, such that the displacement strand is released into the liquid.

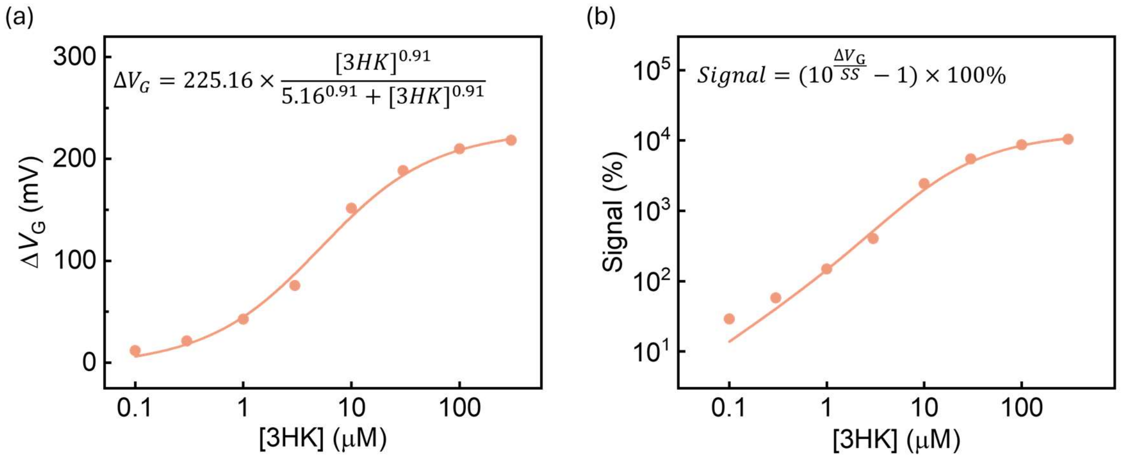


**Figure S6. Binding curve fitting with Hill equation.** (**a**) We performed the binding curve fitting of our CAFET sensor using the $\Delta V_G$ data as a function of analyte concentration. $\Delta V_G$ is proportional to the number of the analyte-bond displacement-strand aptamers. We generated the fitting curve (solid line) using the Hill equation:

$$\Delta V_G = \Delta V_{G\_max} \times \frac{T^n}{T^n + {K_D}^n}$$

where $\Delta V_G$ and $T$ represent the gate voltage shift at various target concentrations and $\Delta V_{G\text{-}max}$, $K_D$, and $n$ are parameters extracted from the fitting. We observed a good fit across the entire concentration range, and extracted the corresponding parameters as shown in the panel. (**b**) We converted the $\Delta V_G$ data points and fitting curve in **a** to the signal in terms of percent FET current change ($\Delta I/I_0$). Considering the current's exponential dependance of $\Delta V_G$, the signal can be converted using this equation:

$$Signal = \frac{\Delta I}{I_0} \times 100\% = \frac{I_0 \times 10^{\Delta V_G/SS} - I_0}{I_0} \times 100\% = (10^{\frac{\Delta V_G}{SS}} - 1) \times 100\%$$

where $SS$ = 108 mV/dec is the slope extracted from the FET transfer curve. We used the data points and fitting curve in **b** to generate **Figure 2c**. The signal conversion from $\Delta V_G$ to signal is not homogeneous across the entire concentration range, which led to the shape deviation of the curve in **b** relative to the fitting in **a**.

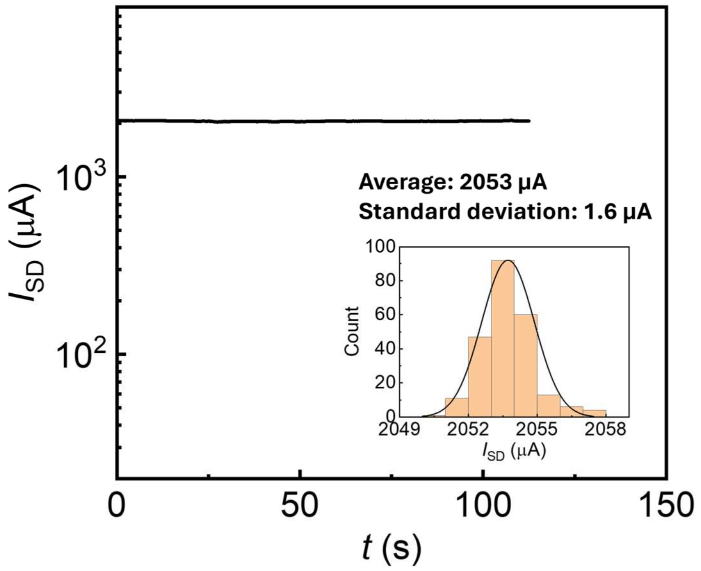


**Figure S7. Standard deviation of the baseline.** We measured the real-time baseline current fluctuation and generated the current histogram in the inset. The standard deviation is roughly 0.08% of the baseline average.

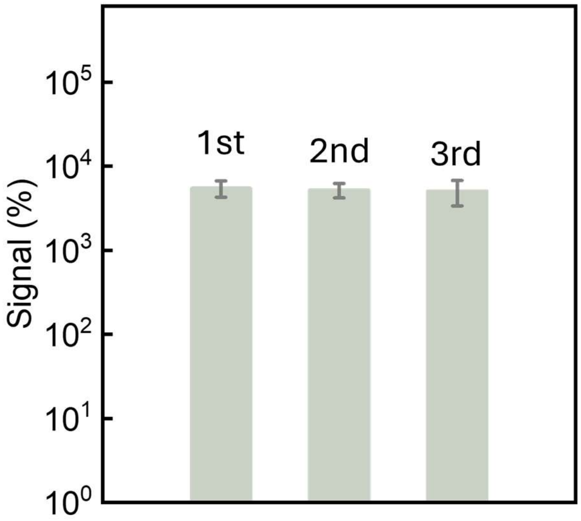


**Figure S8. Reusability of our FET-aptamer sensors.** We measured the baseline response from the starting sensor (1st) and after two rounds of target incubation and regeneration (2nd and 3rd). The signal after re-hybridizing the DS to the aptamer is very close to the signal prior to initial use. Signals were measured in high salt buffer. Error bars were obtained from measurements of three independent sensors.

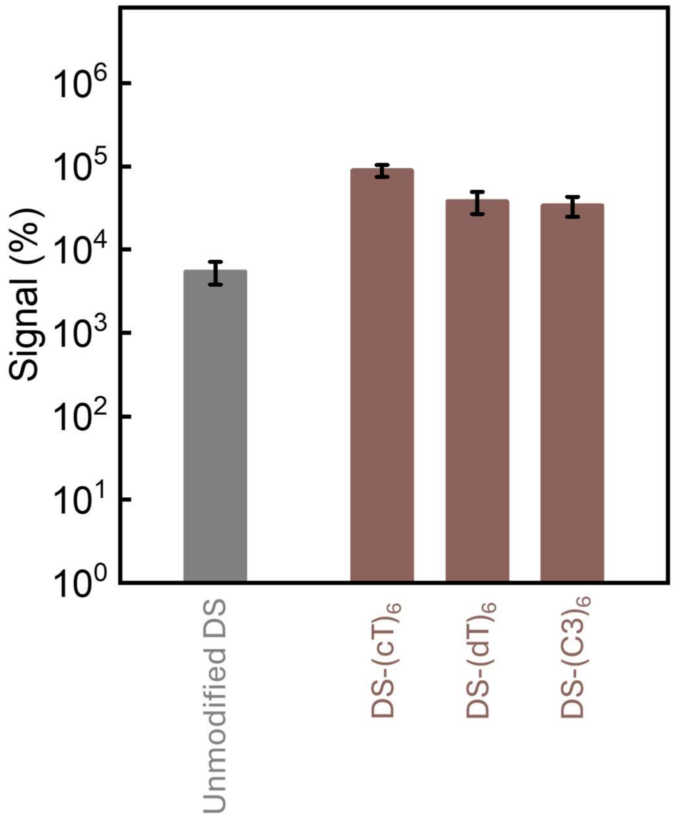


**Figure S9. Comparison of different charge-labeling strategies.** FET-aptamer sensor signal in response to 30 µM 3HK with either unmodified DS or DS labeled with 6-mer chains of oligo-carboxy-dT (cT), oligo-dT, and spacer C3. The signal gain with DS-(dT)$_6$ and DS-(C3)$_6$ is lower than with DS-(cT)$_6$, since each unit of the former contributes one elementary charge while the latter contributes two charges. The sensor was functionalized with 3HK aptamer, and measurements were performed in high-salt buffer. Error bars were obtained from measurements of three independent sensors.

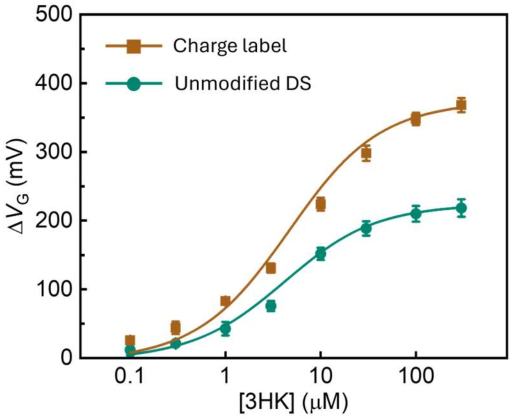


$$\Delta V_G = 373.13 \times \frac{[3HK]^{0.88}}{5.61^{0.88} + [3HK]^{0.88}}$$

$$\Delta V_G = 225.16 \times \frac{[3HK]^{0.91}}{5.16^{0.91} + [3HK]^{0.91}}$$

**Figure S10**. **Aptamer density estimation for our 3HK sensors**. We estimated the aptamer density based on the saturated signal gain $\Delta V_{\text{G-gain}}$ of the charge-labeled DS sequence compared to the unmodified DS. The $\Delta V_G$ data fits well with the Hill equation, from which the saturated $\Delta V_G$ of the two sensors can be extracted (labeled: 373.13 mV; unmodified: 225.16 mV). The measured $\Delta V_{\text{G-gain}}$ = 373.13 mV−225.16 mV = 147.97 mV, which we ascribed to the charge label $(cT)_6$, which contributes 12 elementary charges per label molecule. Since each label molecule binds one aptamer molecule, their molecular density is equal. Therefore, the saturated signal gain can be written as:

$$\Delta V_{\text{G-gai}} = \frac{qN_{\text{charge}}}{C_{\text{DL}}} = \frac{12qN_{\text{label}}}{C_{\text{DL}}} = \frac{12qN_{\text{aptamer}}}{C_{\text{DL}}},$$

where $q$ is the elementary charge and $C_{\text{DL}}$ is the capacitance of the electrical double layer. $C_{\text{DL}}$ is close to 10 μF/cm$^2$ at physiologically high salt strength (S1). Consequently, the aptamer density can be estimated as $N_{\text{aptamer}} = \frac{\Delta V_{\text{G-gai}}\ C_{\text{DL}}}{12q} = \frac{147.97\ \text{mV} \times 10\mu\text{F/cm}^2}{12 \times 1.6 \times 10^{-19}\text{C}} = 8 \times 10^{11}\text{cm}^{-2}$.

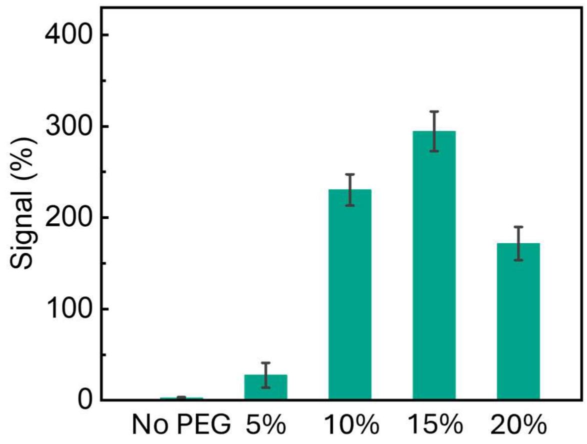


**Figure S11. Biofouling prevention using a PEG2000 passivation layer.** Signal response to 30 μM 3HK from FET-aptamer sensors passivated with various molar fractions of PEG2000. Error bars were obtained from measurements of three independent sensors.

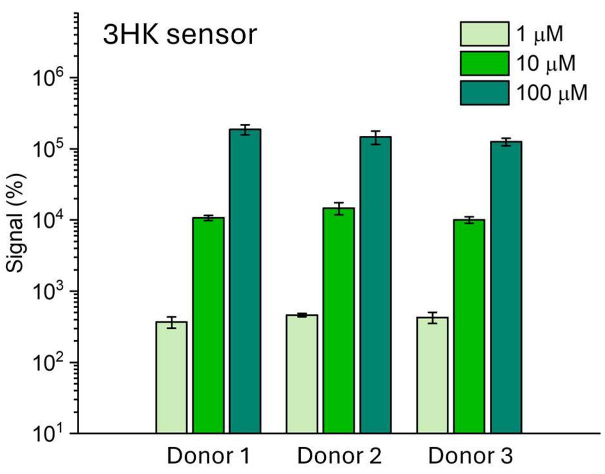


**Figure S12. 3HK sensing in healthy donor plasma samples.** The 3HK sensor was hybridized to a $(cT)_6$-labeled DS. Error bars were obtained from measurements of three independent sensors.

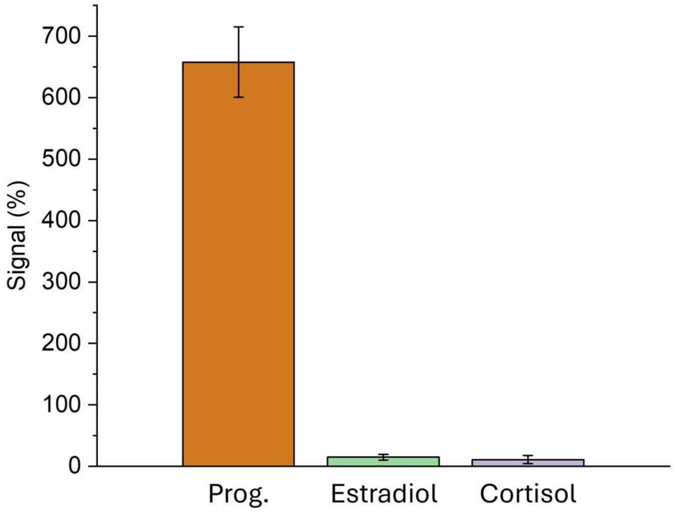


**Figure S13. Selectivity tests of progesterone sensors.** The progesterone sensor was hybridized to unmodified DS and evaluated against a 10 µM concentration of each target in high-salt buffer solution. Error bars were obtained from measurements of three independent sensors.

**Table S1. Sequences of 3HK and progesterone aptamers and their displacement strands.**

| | **Sequence (5'-3')** |
|---|---|
| 3HK aptamer | GGCTCTCGGGACGACACGGGAAGCTTTAGGTTGAGCCATGTGCAG GTCGTCCCTG |
| 3HK aptamer DS | GTCGTCCCGAGAGCC |
| Progesterone aptamer | CGACTGGCAGGGCAGTTAGCCCGGGAACACGATTTTATTGCATGG CTGCCCTGTG |
| Progesterone aptamer DS | CTGCCCTGCCAGTCG |

**Table S2. Benchmark of our sensors against previously reported FET-aptamer sensors.**

| | Analyte | Media | $K_D$ | Current signal at $K_D$ (%) | $\Delta V_G$ at $K_D$ (mV) |
|---|---|---|---|---|---|
| This work | 3HK | High salt buffer | 5.61 µM | **5,238.9%** | **186.5 mV** |
| This work | 3HK | Undiluted plasma | 7.54 µM | **327.3%** | **68.1 mV** |
| This work | Progesterone | High salt buffer | 1.22 µM | **575.5%** | **89.6 mV** |
| This work | Progesterone | Undiluted plasma | 1.78 µM | **107.46%** | **34.2 mV** |
| Ref. S2 | Dopamine | Artificial cerebrospinal fluid | 10 pM | 6.7% | 75 mV |
| Ref. S2 | Serotonin | Artificial cerebrospinal fluid | 10 fM | 14.6% | 8 mV |
| Ref. S2 | Glucose | Ringer's buffer | 100 pM | 25.0% | 15 mV |
| Ref. S2 | S1P | HEPES buffer | 300 pM | 6.4% | 10 mV |
| Ref. S3 | Phenylalanine | Ringer's buffer | 25 nM | 44.4% | 27 mV |
| Ref. S4 | Serotonin | Phosphate buffer | 20 fM | 2.4% | 7 mV |
| Ref. S5 | Cortisol | Artificial sweat | 30 pM | 5.8% | 12 mV |
| Ref. S6 | Dopamine | Artificial cerebrospinal fluid | 6 nM | 12.0% | NA |
